# The Sde phosphoribosyl-linked ubiquitin transferases exploit reticulons to protect the integrity of the *Legionella*-containing vacuole

**DOI:** 10.1101/2023.06.27.546723

**Authors:** Mengyun Zhang, Seongok Kim, Ralph R. Isberg

## Abstract

*Legionella pneumophila* grows within host cells by forming a specialized membrane-bound compartment via the Icm/Dot type IV secretion system (T4SS). T4SS translocated Sde proteins promote phosphoribosyl-linked ubiquitination (pR-Ub) of several mammalian cell proteins, including Rtn4. In response to modification, Rtn4 forms tubular ER aggregates around the *Legionella-*containing vacuole (LCV). The loss of *sde* together with *sdhA* results in severe vacuole disruption at early infection timepoints. We tested if Rtn4 modification allowed it to serve as a physical barrier to protect its degradation from host-derived disruptive compartments. The challenge of the *rtn4*^*-/-*^ macrophages with Δ*sdhA* partially phenocopied the challenge of *rtn4*^*+/+*^ with Δ*sde*Δ*sdhA*, indicating that Rtn4 plays a role in protecting LCV integrity. Depletion of *rtn1* in *rtn4*^*-/-*^ macrophages potentiated vacuole permeability when compared to *rtn4*^*-/-*^ macrophages, consistent with Sde targeting multiple reticulon family members to support vacuole integrity. These results indicate that *L. pneumophila* exploits Rtn4 in cooperation with Rtn1 to establish a replication niche by promoting Sde-mediated tubular ER aggregates, arguing that these proteins function as a physical barrier during early steps of LCV biogenesis.

## Introduction

*Legionella pneumophila* is an intravacuolar pathogen of both amoebae and human hosts (1). As an agent of Legionnaires’ disease, infection is initiated by inhalation of aerosolized contaminated water, followed by bacterial growth within alveolar macrophages (2, 3). Upon internalization, *L. pneumophila* is enclosed in host-derived membrane compartment and secretes over 300 effector proteins via the Icm/Dot type IV secretion system (T4SS) into host cells (4-8). Icm/Dot translocated substrates (IDTS) co-opt host-membrane trafficking pathways to establish a replication-permissive *Legionella*-containing vacuole (LCV) (9). Previous studies demonstrated that LCVs are decorated with endoplasmic reticulum (ER)-associated markers which are tightly linked to supporting the ability of the LCV to bypass degradative host compartments (10-13).

In eukaryotic cells, peripheral ER contains tubular ER networks (polygonal network of tubules) formed by interactions between reticulons (Rtns), DP1/Yop1p and atlastins (14, 15). Evolutionarily conserved reticulon proteins consist of four subfamilies in mammalian cells (Rtns1-4) that are involved in ER-shaping (16, 17). These proteins are exclusively localized to the tubular ER and are involved in stabilizing the high curvature of ER tubules through their two hydrophobic transmembrane hairpins inserted into the lipid bilayer by forming homo-or hetero-Rtn oligomers (14, 18, 19).

Successful intracellular growth of *L. pneumophila* within host cells is dependent on maintaining LCV integrity (20). The bacterial SdhA protein has been demonstrated to play a critical role in preventing the disintegration of the LCV membrane, primarily by preventing attack by disruptive host disruptive compartments derived from the early and recycling endosomes (20, 21). In the absence of *sdhA, L. pneumophila* is exposed to the host cytosol due to compromised vacuole membrane integrity, followed by bacterial degradation by interferon-regulated proteins, which leads to pyroptotic cell death (20, 22).

Sde proteins (SdeABC and SidE) comprise a family of IDTS that each contain an N-terminal deubiquitinase (DUB), a nucleotidase/phosphohydrolase (NP) domain, and a mono-ADP ribosyltransferase (mART) domain. The mART domain ADP-ribosylates host ubiquitin (Ub), serving as a substrate for the NP domain to either hydrolyze the modification or promote transfer of phosphoribosyl (pR)-Ub to either Ser or Tyr residue of host target proteins (23-31). Rtn4 is one of the most efficiently modified targets known (26, 29, 32, 33). Phosphoribosyl-linked Ub of Rtn4 generates a thick tubular Rtn4 aggregate about the LCV soon after bacterial contact with host cells (29). This structure may be important for maintaining vacuole integrity and protecting against host endosomal attack, because LCV disruption occurs rapidly in mutants lacking both *sde* and *sdhA*. This is in contrast to strains lacking only *sdhA*, which show vacuole disruption hours later than the double mutant. That Sde proteins contribute to maintaining vacuole integrity, and an Rtn4-rich tubular structure results from pR-Ub modification, indicates that pR-Ub modified Rtn4 may stabilize the LCV by forming a physical barrier that walls off the LCV, blocking access of host endosomal compartments at early stage of infection (34).

In this work, we determined if Rtn4 plays a role in maintaining vacuole integrity of LCVs. In the course of this study, we determined that multiple reticulon isoforms can support replication vacuole integrity, consistent with pR-Ub modification of multiple reticulon family members driving the formation of physical barrier outside the LCV that blocks access of destabilizing host cell compartments.

## Results and Discussion

### Rtn4 is required for stabilizing LCV integrity

We previously demonstrated that Sde proteins contribute to maintaining the integrity of the *Legionella-*containing vacuole and prevent attack by early endosomal network proteins that act to destabilize the vacuole. Sde proteins are known to promote phosphoribosyl-linked ubiquitination (pR-Ub) of target proteins, with reticulon 4 (Rtn4) being one of the major targets. Furthermore, in response to the presence of Sde, Rtn4-rich tubular ER aggregates are generated surrounding the LCV as a consequence of phosphoribosyl-linked ubiquitination of Rtn4 (34). Sde proteins can target several host proteins other than Rtn4, so there is no clear demonstration that Sde protects from vacuole disintegration by directly targeting Rtn4, although aggregation of the protein about the LCV makes it a likely strategy for protecting against attack by the endosomal network (32). Therefore, we sought to determine if Rtn4 is directly involved in promoting LCV integrity.

To test if Rtn4 plays a role in preventing degradation of the LCV, we took advantage of the fact that LCVs harboring the Δ*sde*Δ*sdhA* mutant are permeable 2 hours postinfection, whereas LCVs harboring single Δ*sdhA* mutants are largely intact (34). We then tested if mutational loss of Rtn4 could phenocopy the Δ*sde* mutation at 2 hpi by measuring vacuole disruption in mouse bone marrow-derived macrophages (BMDMs) from either C57BL/6J (WT) or the congenic *rtn4*^-/-^ lines after challenge with the *L. pneumophila* Δ*sdhA* mutant. LCV integrity was measured by fixing BMDMs at this timepoint, and challenging with anti-*L. pneumophila* to measure accessible bacteria in the presence or absence of chemical permeabilization (Fig. 1A) (20).

**Fig. 1.**
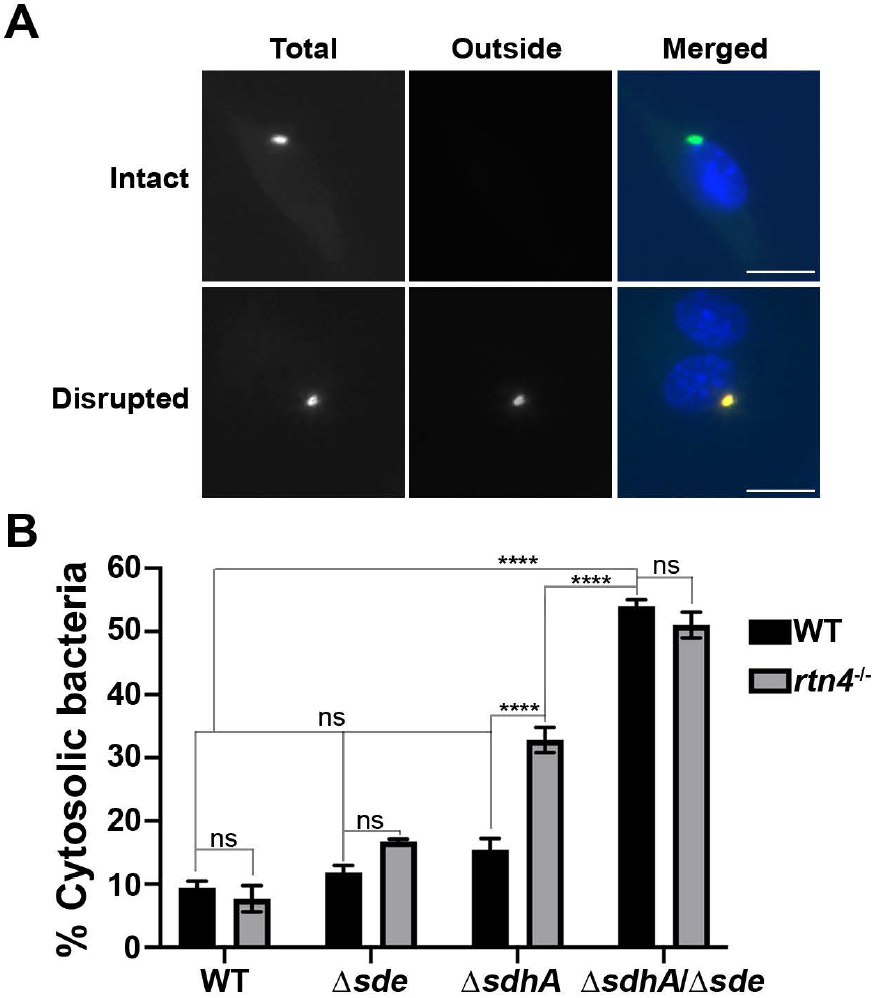
Disintegration of vacuoles harboring the Δ*sdhA* strain is exacerbated by the absence of Rtn4. (A) Examples of cytosolic and vacuolar bacteria. WT or *rtn4*^-/-^ BMDMs were challenged with the noted *Legionella* strains for 2 hr, fixed, probed with anti-*L. pneumophila* (Alexa Fluor 594 secondary, red), permeabilized (Materials and Methods), and reprobed with anti-*L. pneumophila* (Alexa Fluor 488 secondary, green). Cytosolic bacteria are shown in yellow in the merged image and vacuolar bacteria are shown in green. The scale bar represents 10 μm. (B) The vacuole integrity of indicated strains in WT or *rtn4*^-/-^ BMDMs at 2 hr post-infection. The percentage of vacuole disruption was calculated as the following: the bacteria (positively stained without permeabilization) / total bacteria (positively stained with permeabilization). Data are shown as mean ± SEM; three biological replicates with 50 LCVs counted per replicate. Statistical analysis was performed using two-way ANOVA with Tukey’s multiple comparisons, with significance represented as: ns, non-significant; ****p < 0.0001.

The absence of Rtn4 in BMDMs had no effect on the levels of permeability of LCVs to antibody staining after challenge with either a WT or Δ*sde* strains (Fig. 1B, compare WT to *rtn4*^*-/-*^ (34)). In contrast, challenge of *rtn4*^*-/-*^ macrophages with Δ*sdhA* strains resulted in a significant increase of disrupted vacuoles when compared to WT macrophages, with loss of Rtn4 resulting approximately 2X more antibody-accessible vacuoles (Fig. 1B; Δ*sdhA* in WT BMDM vs Δ*sdhA* in *rtn4*^-/-^ BMDM). This result is consistent with pR-Ub modified Rtn4 playing a role in supporting vacuole integrity. Interestingly, vacuole disruption after challenge of Δ*sdhA* strain in *rtn4*^-/-^ BMDMs did not result in an exact phenocopy of the Δ*sdhA*Δ*sde* strain, as the latter showed a significantly higher number of degraded vacuoles after challenge of BMDMs from either mouse strain (Fig. 1B; compare Δ*sdhA* in *rtn4*^*-/-*^ to Δ*sdhA*Δ*sde* in *rtn4*^*+/+*^). As other reticulon family members are known to be targeted for Sde modification (32), we tested the model that multiple reticulon family members collaborate to maintain vacuole integrity after challenge of macrophages with the Δ*sdhA* strain.

### *L. pneumophila* exploits Rtn1 in cooperation with Rtn4 to maintain vacuole integrity

Reticulon 1 (Rtn1) is an abundant reticulon family member that has been shown to be targeted by Sde proteins for pR-Ub modification based on mass spectrometry analysis (32). Therefore, we determined if modification of Rtn1 played a role in maintaining the integrity of the LCV in cooperation with Rtn4. To this end, Rtn1 was depleted by siRNA in *rtn4*^*+/+*^ or *rtn4*^-/-^ BMDMs prior to bacterial challenge and vacuole integrity was measured at 2 hpi. Depletion of *rtn1* with siRNA resulted in approximately 75% loss of transcription and approximately 50% loss of steady-state levels of RTN1 protein (Fig. 2A). Under these conditions, depletion of *rtn1* in the *rtn4*^*-/-*^ BMDMs resulted in increased vacuole disruption after challenge with *L. pneumophila* Δ*sdhA*, with 38% higher vacuole permeability as compared to the scrambled control in the *rtn4*^*-/-*^ cells (Fig. 2B; compare si-NT to si-Rtn1 in Δ*sdhA rtn4*^-/-^; p < 0.0001, two-way ANOVA with Tukey’s multiple comparisons). As a consequence of loss/depletion of two reticulon isoforms, approximately 50% of the LCVs were no longer intact (Fig. 2B), consistent with these proteins collaborating to support LCV integrity.

**Fig. 2.**
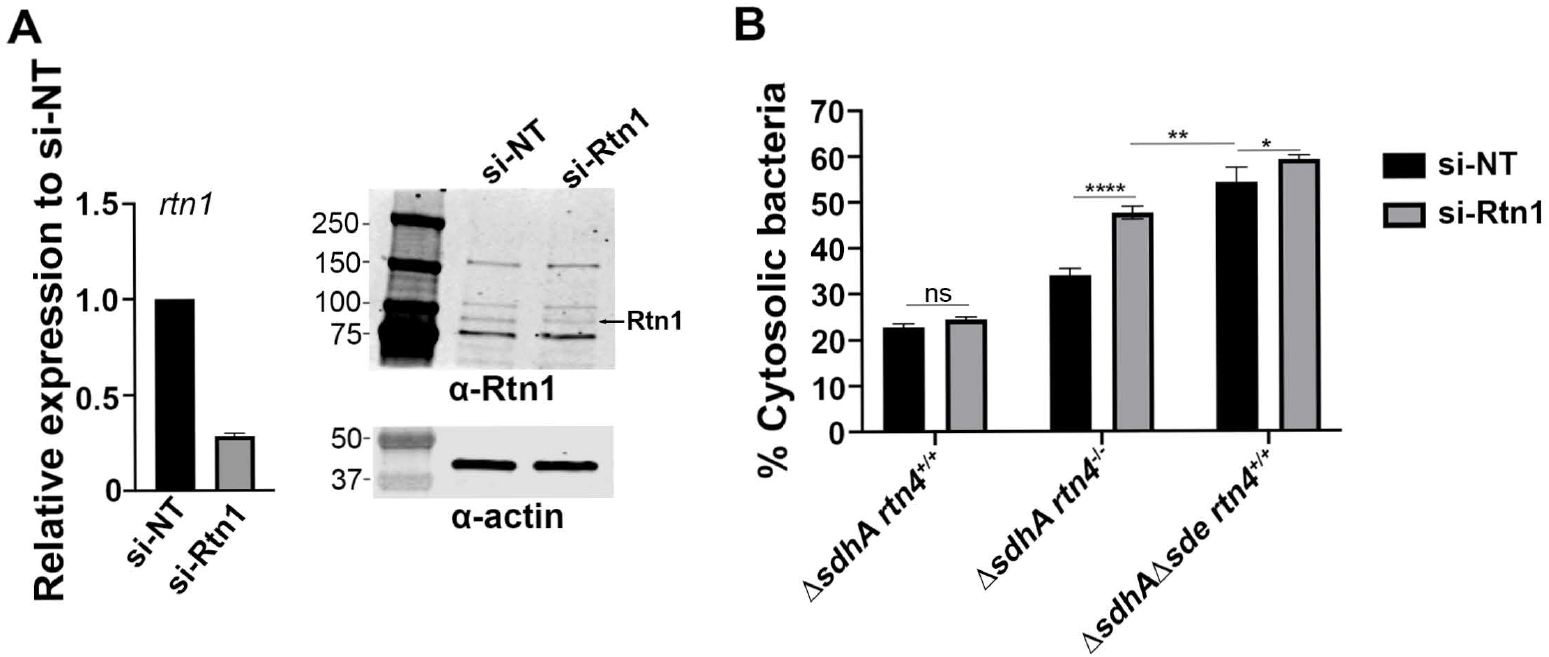
Rtn1 knockdown in *rtn4*^*-/-*^ BMDMs aggravates disruption of LCVs harboring the Δ*sdhA* strain. (A) Depletion efficiency of si-Rtn1 in *rtn4*^*-/-*^ BMDMs. Left: transcript abundance of *rtn1* relative to si-NT. Right: immunoblot of SDS-polyacrylamide gel probed with anti-Rtn1 after depletion with si-NT or si-Rtn1. β-actin was used as loading control. (B) Effect of si-Rtn1 on vacuole integrity at 2 hr post-infection. WT or *rtn4*^*-/-*^ BMDMs were pretreated with si-NT (non-targeting) or si-Rtn1 and challenged with the noted *L. pneumophila* strains for 2 hrs, followed by probing as described (Materials and Methods). Data shown as mean ± SEM for three biological replicates. Statistical significance was tested using two-way ANOVA with Tukey’s multiple comparisons; ns (non-significant), *p < 0.05, **p < 0.01, ****p < 0.0001. At least 50 LCVs were counted per replicate.

In this report we provide evidence that Sde modification of multiple reticulon isoforms is a key step in supporting LCV integrity, as revealed by identifying determinants necessary for vacuole stability in the absence of SdhA function. The previously demonstrated ability of Sde proteins to support LCV integrity raised the possibility that modification of reticulons was a key step in vacuole protection. Evidence is provided here that manipulation of reticulon dynamics by *L. pneumophila* is indeed central to this process, as loss of Rtn4 results in a marked depression in the ability of Sde to support vacuole integrity (Fig. 1). Depletion of Rtn1 by siRNA aggravated the defect observed in *rtn4*^-/-^ cells, indicating that multiple reticulon family members (each of which are Sde targets) contribute to this role. It should be pointed out that the depletion of Rtn1 in the *rtn4*^-/-^ BMDMs does not provide an equivalent phenotype to the Δ*sdhA*Δ*sde* strain (Fig. 2; compare si-Rtn1 in Δ*sdhA rtn4*^-/-^ to si-NT in Δ*sdhA*Δ*sde rtn4*^-/-^). Other reticulon proteins such as Rtn2 or Rtn3, may be similarly involved in protecting LCV integrity, or the siRNA directed against Rtn1 may not be sufficient to give full penetrance of the phenotype. In either case, the results presented here are consistent with Sde-modifed reticulons protecting from endosomal attack. As was hypothesized in our previous communication, aggregation of Rtn1/Rtn4 is likely to provide a physical barrier that prevents access of these endosomal components to the LCV. The fact that loss of reticulon function prevents Sde from protecting against LCV disintegration is an important result that strongly supports the concept of a barrier surrounding the LCV.

## Materials and Methods

### Bacterial strains, cell culture and media

*L. pneumophila* strains were propagated in liquid N-(2-acetamido)-2-aminoethanesulfonic acid or on solid charcoal buffered yeast extract (CYE) media containing 0.4g/l iron (III) nitrate, 0.135 g/ml cysteine, and 1% α-ketoglutaric acid. 40μg/ml kanamycin and 5% (vol/vol) sucrose were added when appropriate. Primary bone marrow-derived macrophages (BMDM) from the femurs and tibias of female C57BL/6J or congenic RTN4^-/-^ (kind gift from Dr. William Sessa) were prepared and cultured as described (11, 34).

For construction of *L. pneumophila* Δ*sdhA*Δ*sde* lacking flagellin, *flaA* gene was deleted in the Δ*sdhA*Δ*sde* strain (34) by allielic exchange recombination using suicide plasmid pSR47s as previously described (35). All strains used in this study were Δ*flaA*.

### Assay for LCV integrity

To test vacuole integrity, C57BL/6 (WT) or congenic *rtn4*^*-/-*^ BMDMs were challenged with the noted *L. pneumophila* strains at an MOI = 1, contact was initated at 400 x g for 5 min in a table top centrifuge and incubated at 37°C, 5% CO_2_. At 1 hr post-infection (hpi), the macrophages were rinsed with 3X PBS, replenished with fresh medium followed by 1hr additional incubation at 37°C, 5% CO_2_. At 2 hpi, infection mixtures were washed 3x in PBS at room temperature, fixed in PBS containing 4% paraformaldehyde, then probed with mouse anti-*L. pneumophila* (Bio-Rad, Cat# 5625-0066, 1:10,000) followed by secondary probing with goat anti-mouse Alexa Fluor 594 (Invitrogen, Cat# A11005, 1:500) to identify disrupted vacuoles as described (20). After washing 3X with PBS, LCVs were permeabilized by 5 min incubation with -20°C methanol, washed 3X in PBS and blocked with PBS containing 4% BSA for 30 min prior to a second probing with mouse anti-*L. pneumophila* (Bio-Rad, Cat# 5625-0066, 1:10,000). All bacteria (both intact and disrupted vacuoles) were identified by goat anti-mouse IgG Alexa Fluor 488 (Invitrogen, Cat# A11001, 1:500). Images were taken using Zeiss observer Z1at 63X and individual BMDMs were scored for disruption based on staining with goat anti-mouse Alexa Fluor 594 (antibody accessible in the absence of methanol) as described previously (34).

### RNA interference

5 × 10^6^ cells of BMDMs were seeded in 10 cm dishes filled with 10 mls RPMI medium containing 10% FBS and 10% supernatant produced by 3T3-macrophage colony stimulating factor (mCSF) cells and incubated overnight. Cells were lifted with ice-cold PBS, resuspended in the RPMI medium, aliquoted into 1.5 ml microcentrifuge tubes containing 1 x 10^6^ cells, followed by centrifugation at 200 x g for 10 min. The cell pellets were resuspended in nucleofector buffer (Amaxa Mouse Macrophage Nucleofector Kit, Cat# VPA-1009) and 2 µg of siRNA was added (siGENOME smart pool, Dharmacon). Cells were transferred to a cuvette and nucleofected in the Nucleofector 2b Device using Y-001 program settings according to manufacturer’s instructions. Nucleofected macrophages were immediately recovered in the medium and plated in 24-well plate containing coverslips at 2 x 10^5^/well for microscopy assay or in 12-well plate at 1 x 10^6^/well for RNA extraction or immunoblotting for 48 hrs.

### Quantitative RT-PCR

BMDM cells were challenged with post-exponentially grown *L. pneumophila* strains at MOI=1, washed at 1 hpi, replenished with fresh medium and incubated for additional 1 hr at 37°C, 5% CO_2_. At 2 hpi, total RNA was extracted using Trizol according to manufacturer’s instructions and cDNA was synthesized using SuperScript IV VILO Master Mix with ezDNase enzyme (Thermo Fisher Scientific, Cat#11766050). PowerUp SYBR Green Master Mix was then used for qRT-PCR reactions using Second Step instrument (ABI). Transcriptional levels of genes were normalized to GAPDH.

### Immunoblotting

The efficiency of siRNA knockdown in nucleofected cells was determined by Western blotting. Macrophages plated in 12-well plate were lysed by incubating NP40 buffer (Thermo Fisher Scientific, Cat# J60766.AP) on ice for 30 min and protein concentration was measured by Bradford assay. Proteins in SDS-PAGE sample buffer were boiled for 10 min, fractionated by SDS-PAGE and transferred to nitrocellulose membranes. The membrane was blocked in 50 mM Tris-buffered saline/0.05% Tween 20 (TBST, pH 8.0) containing 4% nonfat milk (blocking buffer) for 1 hr at room temperature and incubated with anti-RTN1 (MyBioSource, Cat#MBS9200801) or β-actin (Invitrogen, Cat# PA1-183, 1:1,000) in blocking buffer at 4°C overnight. The membranes were washed 3X with TBST and incubated with secondary antibody (Li-Cor Biosciences, Cat#926-32211, 1:20,000) in blocking buffer for 1 hr at room temperature. Images were captured and analyzed using Odyssey Scanner and the image Studio software (LI-COR Biosciences).

